# Kin selection and sexual conflict: male relatedness and familiarity do not affect female fitness in seed beetles

**DOI:** 10.1101/637926

**Authors:** Elena C. Berg, Martin I. Lind, Shannon Monahan, Sophie Bricout, Alexei A. Maklakov

**Affiliations:** Department of Computer Science, Mathematics, and Environmental Science, The American University of Paris, France; Department of Ecology and Genetics, Animal Ecology, Uppsala University, Uppsala, Sweden; School of Biological Sciences, University of East Anglia, Norwich Research Park, Norwich, UK

**Keywords:** sexual conflict, kin selection, mate harm, *Callosobruchus maculatus*

## Abstract

Theory maintains that kin selection can mediate sexual conflict because within-group male relatedness should reduce male-male competition, thereby reducing collateral harm to females. We tested whether male relatedness and familiarity can lessen female harm in the seed beetle *Callosobruchus maculatus.* Neither male relatedness nor familiarity influenced female lifetime reproductive success or individual fitness. However, male relatedness, but not familiarity, marginally improved female survival. Because male relatedness improved female survival in late life when *C. maculatus* females are no longer producing offspring, our results do not provide support for the role of kin selection in mediating sexual conflict. The fact that male relatedness improves the post-reproductive part of female life cycle strongly suggests that the effect is non-adaptive. We discuss adaptive and non-adaptive mechanisms that could result in reduced female harm in this and previous studies and suggest that cognitive error is a likely explanation.

## 1. INTRODUCTION

Males and females have different routes to successful reproduction [1], and this can lead to evolutionary conflict between the sexes [2–4]. One extreme form of this conflict is mate harm, when one sex (usually the male) physically injures the opposite sex (usually the female) [4, 5]. Male mate harm occurs in many animals and has been especially well studied in insects, including the bed bug *Cimex lectularius,* the seed beetle *Callosobruchus maculatus,* and the fruit fly *Drosophila melanogaster.* Male bed bugs stab females with their genitalia, inseminating females directly into the abdominal cavity [6, 7]. Male seed beetles have spines on their intromittent organs that pierce holes in the female’s genital tract, reducing female longevity [8–10]. In the fruit fly, males may harass females during courtship [11–14] or physically harm them during the mating process [11, 15]. *Drosophila* and *C. maculatus* ejaculates also contain accessory gland proteins that modulate female reproductive behaviour [5, 16–18]. Mate harm can evolve for two reasons. One possibility is that the harm itself increases male fitness by causing females to allocate more resources into current reproduction and away from future reproductive attempts with other males [19]. A better-supported alternative in most cases is that mate harm is simply a deleterious side effect of male-male competition over fertilization [4, 20, 21].

Kin selection theory [22, 23] suggests that the level of genetic relatedness among competing males can moderate sexual conflict in viscous populations [24–27]. In populations in which close adult relatives are likely to interact, males would gain indirect fitness benefits by helping their close male kin to reproduce. Such cooperation might take the form of reducing mate harm to facilitate sexual access to those females. As Chippindale *et al.* [28] point out, for this kind of kin selection to occur, three conditions must be met: 1) males must harm their mates in some way, thereby reducing female reproductive success; 2) there must be some mechanism in place for reliably recognizing kin; and 3) groups of related males must have a reasonable chance of encountering each other during the reproductive period.

A series of recent studies on *Drosophila melanogaster* has investigated this possible role of kin selection in moderating mate harm, with conflicting results [28–33]. In support of the kin selection hypothesis, Carazo *et al.* [28] found that females exposed to groups of three full-sib brothers did indeed have higher lifetime reproductive success and slower reproductive ageing than females exposed to trios of unrelated males. Groups of brothers were less aggressive towards each other, courted females less vigorously, and lived longer. Finally, Carazo *et al.* [28] found that when two brothers were housed with an unrelated male, that unrelated male sired more than one-third of the offspring, suggesting that groups of cooperating relatives are vulnerable to invasion by non-cooperative non-relatives.

One important point to consider in the Carazo *et al.* [28] study is that it confounded familiarity and relatedness. Brothers used in the study had also been reared together in the same vial, whereas the unrelated males had been raised in separate vials. Hollis *et al.* [30] conducted a follow-up study on a different population of *Drosophila* in which they controlled for familiarity by testing the effect of brothers raised together versus apart. Females exposed to brothers raised together had higher lifetime reproductive success, but this effect disappeared when females were raised with brothers raised apart. Hollis *et al.* [30] concluded that familiarity and not relatedness *per se* was likely driving the patterns Carazo *et al.* [28] observed.

However, one weakness of Hollis *et al.’s* [30] study is that they did not include a treatment with unrelated males that had been reared together. Without this treatment it is impossible to conclude whether familiarity alone would be sufficient for cooperation. To address this issue, Chippindale *et al.* [29] performed a fully crossed experiment in which they exposed females to brothers that had been raised together, brothers that had been raised apart, and unrelated males that had been raised apart. Unlike Carazo *et al.* [28], they found no evidence that either familiarity or relatedness among males had any effect on female lifespan or reproductive success, a result corroborated in a separate study by Martin and Long [31]. Further, Chippindale *et al.* [29] found no evidence that unrelated or unfamiliar males sired a disproportionate number of offspring when introduced to pairs of brothers or males raised in the same environment. They conclude that cooperation among males does not appear to lower mate harm in this system, at least not in the populations they examined. Finally, a later study by Le Page *et al.* [32] suggested that both relatedness and familiarity are required for reduced female harm in *D. melanogaster* in the population used by Carazo *et al.* [28, 33].

These conflicting results suggest that, at least in *Drosophila,* it remains unclear what role kin selection plays in mediating male-male cooperation and mate harm. Moreover, if we are to understand whether inclusive fitness benefits mediate sexual conflict in the animal kingdom, we need to expand our research focus into other model systems. One excellent candidate is the seed beetle *C. maculatus.* As described above, male seed beetles inflict physical harm on their mates [9, 10] and this species has been used routinely as a model system to study the economics and genetics of sexual conflict over lifespan and reproduction [34–36].

A recent study by Lymbery *et al.* [37] generally supported the importance of male relatedness in mediating male harm to females. They found that females housed with familiar brothers produced more offspring, suggesting that relatedness and familiarity among males act together to reduce male-induced harm to females. However, the beetles in the Lymbery *et al.* [37] study were provided with Baker’s yeast, which is rather unusual for a species that inhabits human grain storages and is commonly kept in the laboratory as a capital breeder that is aphagous in the adult stage. Furthermore, while *C. maculatus* beetles can technically ingest yeast, yeast consumption *per se* does not necessarily have a positive effect on longevity, fecundity or offspring production [38] suggesting that access to yeast is not a part of a normal life cycle of this species. Therefore, we investigated the effect of male relatedness and familiarity in a large outbred and well-described population of *C. maculatus* (SI USA) that was not provided with yeast in the adult stage, which is in line with the recent evolutionary history of this species and this population. We conducted a fully crossed experiment with respect to male relatedness and familiarity, quantifying the lifetime reproductive success and lifespan of virgin females exposed to four different trios of males: 1) brothers raised together, 2) brothers raised apart, 3) unrelated males raised together, and 4) unrelated males raised apart. If kin selection is indeed mediating sexual conflict in this system, then brothers raised together should exhibit less mate-harming behaviour than other groups, resulting in higher relative fitness of females.

## 2. METHODS

### (a) Study system

Seed beetles are common pest of stored legumes indigenous to Asia and Africa. Females lay their eggs on the surface of dried beans. Once the larvae hatch, they burrow into the bean and eclose as reproductively mature adults approximately 23-27 days later. *C. maculatus* are facultatively aphagous, obtaining all the nutrients they require for survival and reproduction during the larval stage [39]. Adult feeding increases fecundity and longevity [39]. Early studies used a combination of yeast and sugar solutions, so it was difficult to disentangle the effect of the separate components in fitness-related traits. Ursprung *et al.* [38] found that sugar solution and water do increase fecundity and longevity, but there was no effect of yeast on these key life-history traits. Lymbery *et al.* [37] provided their study beetles with *ad lib* access to yeast, but there was no obvious benefit in terms of fecundity or longevity, although the direct comparison is not possible because their study did not include standard aphagous conditions.

The study population was derived from an outbred South Indian stock population (“SI USA”) of *C. maculatus* originally obtained from C. W. Fox at the University of Kentucky, USA, and then subsequently moved to Uppsala University and finally to The American University of Paris three months prior to the first block of the experiment. The original SI USA stock population was collected from infested mung beans *(Vigna radiata)* in Tirunelveli, India in 1979 [40]. Both prior to and during the experiment, beetles were cultured exclusively on mung beans and kept at aphagy (no food or water) in climate chambers at 29°C, 50% relative humidity and a 12:12 h light:dark cycle. One great advantage of this system is that the laboratory conditions closely resemble natural conditions, because these beetles have associated with dried legumes for thousands of years and their life history is adapted to life in a storage environment [41, 42].

### (b) Establishing the four treatment groups

The experiment was carried out in two blocks. The first block was completed in 2015. In 2018 the experiment was replicated and expanded to provide additional data on daily fecundity of females. During both blocks, base populations of beetles were kept in 1L jars with 150g of mung beans, and approximately 250 newly hatched beetles were transferred to new jars with fresh beans every 24 days on a continual basis. From this base population, we established four different treatment groups (with a goal of approximately N = 75 each in each block, 150 total): **1) related familiar (RF), 2) related unfamiliar (RU), 3) non-related familiar (NF)**, and **4) non-related unfamiliar (NU)**. In the RF treatment, three full-sib brothers were housed in a Petri dish together for 24 hours before being added to a dish with a (non-related) female. In the RU treatment, three brothers with no prior experience with each other were added all at once to a dish with a female. In the NF treatment, three non-related males were housed together for 24 hours before being added to a dish with a female. Finally, in the NU treatment, three non-related and unfamiliar males were added to a dish with a female.

To generate full sibling brothers for the RF and RU treatments, we transferred a random subset of beans with developing larvae into virgin chambers (aerated plastic culture plates with a separate well for each individual) and monitored the virgin chambers daily. Approximately one day after hatch, we randomly paired 180 males and females and placed them into 180 60-mm Petri dishes with 75 beans each. We then removed the males and females after 48 hours and allowed the eggs to develop. Since females can lay up to 65 eggs per day (Berg, unpublished data), we wanted to provide enough beans that females would lay only one egg on each bean. Before the offspring hatched, we transferred the fertilized beans from the Petri dishes to virgin chambers, carefully marking which beans came from which parents, and monitoring hatch daily. Once these offspring hatched, we set up the four different treatment groups above.

For both “familiar” treatments, trios of males were introduced to each other on the same day that they hatched. For both “unfamiliar” treatments, males were housed in their separate virgin cells until one day post hatch and then introduced together with the female without any time to acclimatize to each other. In all treatments, females were randomly selected from the base population one day after hatch and were unrelated to the males. Males from the non-related treatments were randomly selected from the base population as well. All males and females used in this study were maintained as virgins prior to the pairing. The Petri dishes in which males and females were housed measured 100-mm and contained 150 beans. This number is sufficient to allow females to lay just one egg per bean, reducing any larval competition that might affect data on reproductive success. All sets of brothers used in this study came from different parents, thus obviating the need to control for parental identity in the analyses.

Three days after the trios of males were introduced to females, we swapped out all the males for new males. This was done to reduce the variance caused by male condition or behaviour on female reproductive success or lifespan. In preparation for this, we set up new trios of freshly hatched related familiar and non-related familiar males one day before. For all the “related” dishes, we used brothers of the previous trio. Since fewer males were eclosing this late in the hatch cycle, we had to use slightly older males in some cases. We excluded few females that escaped/died from unnatural causes resulting in slight deviations from the initial sample size (N = 75 for RF, N = 75 for RU, N = 71 for NF, and N = 76 for NU in the first block; N = 75 for all treatments in the second block).

### (c) Lifespan and fitness assays

During both blocks of the experiment, we conducted both lifespan and fitness assays for each female within each treatment group. For lifespan assays, we monitored the Petri dishes daily and recorded the date of death of each female. Once all adults were dead, we removed them from the dishes. We collected two kinds of fitness data. During the first block of the experiment, we measured total offspring production only. We did this by counting the number of eclosed young per female, a standard measure of lifetime reproductive success in this system. To facilitate the counting of offspring, we froze the dishes 37 days after the initial pairing, well after all the offspring had eclosed but before a subsequent generation could develop.

During the second block, we also measured daily offspring production for each female. To do this we moved the female and males to new Petri dishes with new beans every 24 hours until the female died (maximum of 9 sets of Petri dishes per female). Approximately 37 days later, we froze the dishes and counted number of eclosed offspring per day per female.

### (d) Statistical analyses

Before analysis, we excluded all individuals that did not reproduce (NF = 2, NU = 4, RF = 4, RU = 10). We analysed the lifetime offspring production as well as age-specific reproduction using a generalized mixed effect model with a Poisson error structure implemented in the *lme4* package in *R 3.3.3.*, treating Relatedness and Familiarity as crossed fixed factors. We tested for overdispersion using the *dispersion_glmer* function in the *blmeco* package, and if above 1.4, we controlled for overdispersion by adding a subject-level random effect. For total reproduction, we used Block as a random factor.

Age-specific reproduction and individual fitness was only analysed for Block 2, as this was the only block where age-specific fecundity data was collected. For age-specific reproduction, we included Relatednes and Familiarity as crossed fixed factors, as well as all interactions with Age and Age^2^. In addition, we also included Age at last reproduction (ALR) as a crossed covariate. Age and ALR were scaled and centered before analysis (mean = 0, s.d. = 1) and we used the *bobyqa* optimizer as well as increased the default number of iterations to 10.000 in order to obtain good model convergence. For all mixed-effect models, chi-square tests of fixed effects were performed using the *car* package.

Individual fitness (λ_ind_) was calculated from the life-table of age-specific reproduction [43, 44], with a development time of 23 days, by solving the Euler-Lotka equation for each individual using the *lambda* function in the *popbio* package. λ_ind_ was then analysed in a linear model using Relatedness and Familiarity as crossed fixed factors.

Survival was analysed in a Cox proportional hazard model using the *coxme* package, with Relatedness and Familiarity as crossed fixed factors, and Block as a random effect.

## 3. RESULTS

We measured lifetime reproductive success (total number of eclosed offspring) of individual females introduced to one of four different groups of male trios: brothers raised together (related familiar, or RF, N = 146), brothers raised separately (related unfamiliar, RU, N = 140), non-related males raised together (non-related familiar, NF, N = 144), or non-related males raised separately (non-related unfamiliar, NU, N = 147). There was no significant difference in female lifetime reproductive success between the four treatments (Relatedness: χ^2^ = 1.37, df = 1, p = 0.392; Familiarity: χ^2^ = 0.00, df = 1, p = 0.999; Relatedness × Familiarity: χ^2^ = 0.0015, df = 1, p = 0.969; Figure 1). If anything, mean number of eclosed young was slightly higher for the non-related treatments. If the non-reproducing females are included in the dataset, we actually find higher reproduction in the non-related treatment group (Relatedness: χ^2^ = 4.30, df = 1, p = 0.038; Familiarity: χ^2^ = 1.33, df = 1, p = 0.248; Relatedness × Familiarity: χ^2^ = 0.240, df = 1, p = 0.624, Supplementary figure 1). We did find different shapes of the age-specific fecundity, illustrated by the significant interaction Relatedness × Familiarity × Age^2^ (Table 1, Figure 2). However, we found no effect on individual fitness λind (Relatedness: F = 0. 026, df = 1, p = 0.872, Familiarity: F = 0.244, df = 1, p = 0.622, Relatedness × Familiarity: F = 0.072, df = 1, p = 0.789).

**Figure 1.**
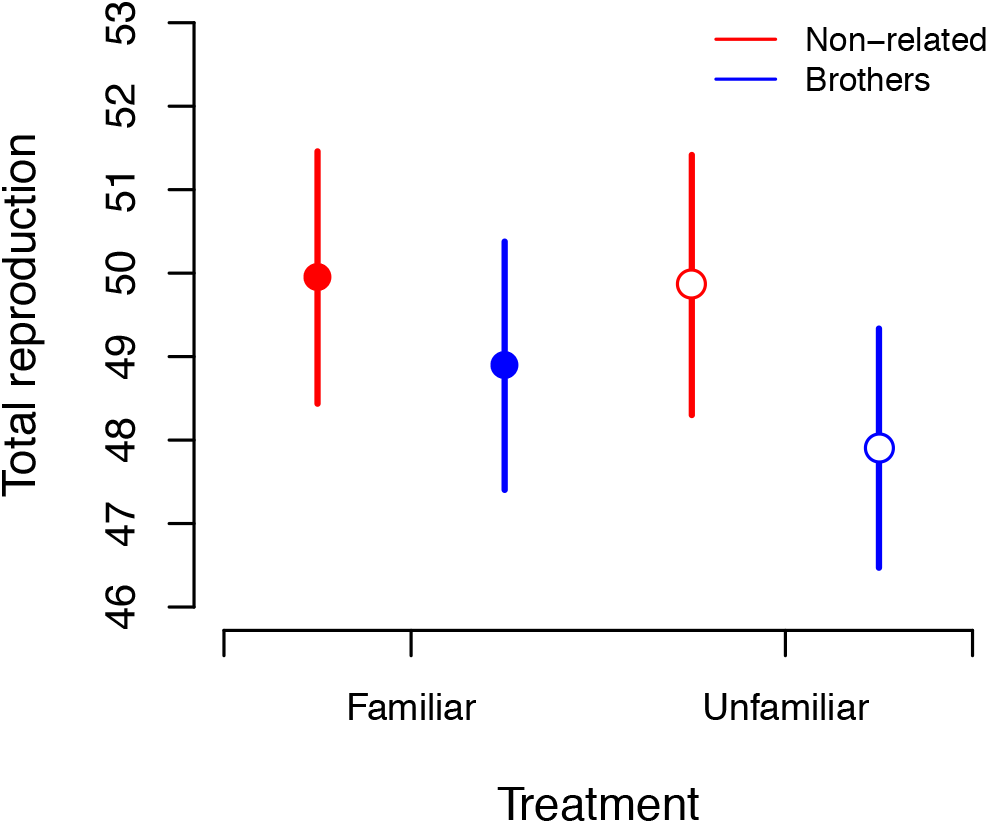
Lifetime reproductive success LRS (number of adult offspring) by treatment group: brothers (blue), non-related males (red), familiar individuals (solid symbols) and unfamiliar individuals (open symbols). Symbols represent mean ± SE.

**Figure 2.**
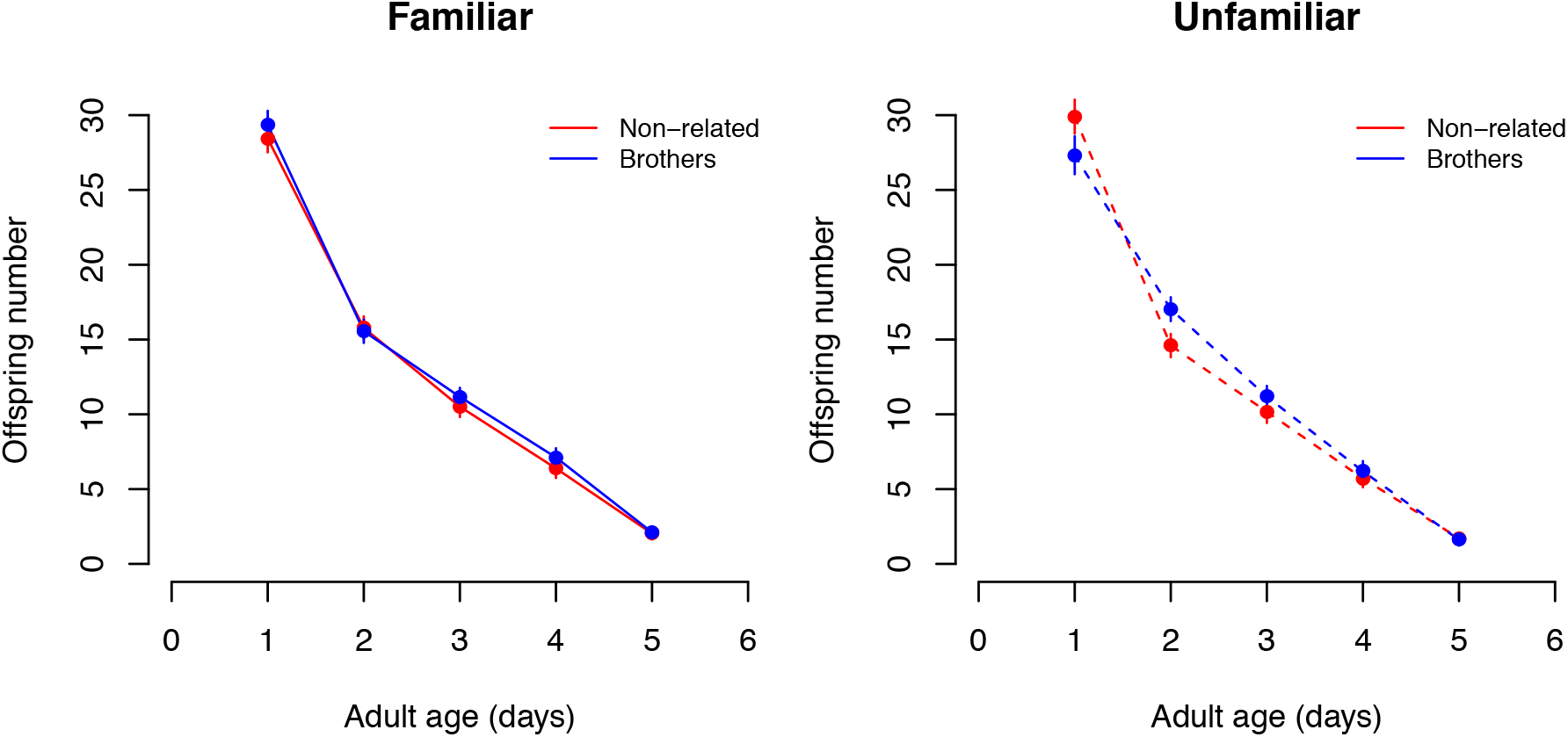
Age-specific reproduction for females mated with (A) familiar and (B) non-familiar males. Brothers are shown as blue, and non-related males as red.

**Table 1.**
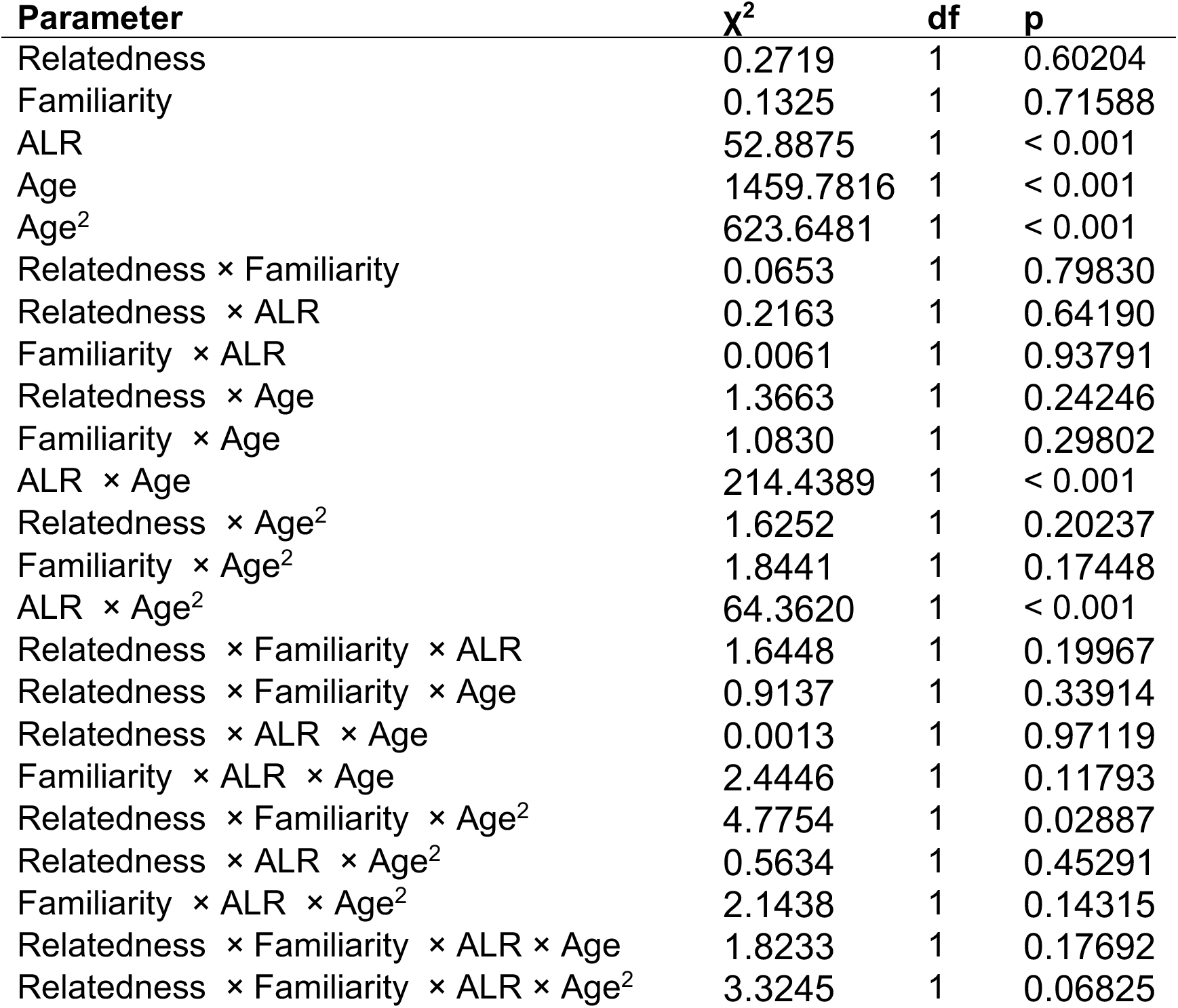
Age-specific reproduction, result from generalized linear model with poisson error structure.

| Parameter | $\chi^2$ | df | p |
| --- | --- | --- | --- |
| Relatedness | 0.2719 | 1 | 0.60204 |
| Familiarity | 0.1325 | 1 | 0.71588 |
| ALR | 52.8875 | 1 | < 0.001 |
| Age | 1459.7816 | 1 | < 0.001 |
| Age<sup>2</sup> | 623.6481 | 1 | < 0.001 |
| Relatedness × Familiarity | 0.0653 | 1 | 0.79830 |
| Relatedness × ALR | 0.2163 | 1 | 0.64190 |
| Familiarity × ALR | 0.0061 | 1 | 0.93791 |
| Relatedness × Age | 1.3663 | 1 | 0.24246 |
| Familiarity × Age | 1.0830 | 1 | 0.29802 |
| ALR × Age | 214.4389 | 1 | < 0.001 |
| Relatedness × Age<sup>2</sup> | 1.6252 | 1 | 0.20237 |
| Familiarity × Age<sup>2</sup> | 1.8441 | 1 | 0.17448 |
| ALR × Age<sup>2</sup> | 64.3620 | 1 | < 0.001 |
| Relatedness × Familiarity × ALR | 1.6448 | 1 | 0.19967 |
| Relatedness × Familiarity × Age | 0.9137 | 1 | 0.33914 |
| Relatedness × ALR × Age | 0.0013 | 1 | 0.97119 |
| Familiarity × ALR × Age | 2.4446 | 1 | 0.11793 |
| Relatedness × Familiarity × Age<sup>2</sup> | 4.7754 | 1 | 0.02887 |
| Relatedness × ALR × Age<sup>2</sup> | 0.5634 | 1 | 0.45291 |
| Familiarity × ALR × Age<sup>2</sup> | 2.1438 | 1 | 0.14315 |
| Relatedness × Familiarity × ALR × Age | 1.8233 | 1 | 0.17692 |
| Relatedness × Familiarity × ALR × Age<sup>2</sup> | 3.3245 | 1 | 0.06825 |

When we measured the lifespan of the females introduced to the different treatment groups, we found that male relatedness, but not familiarity, improved female survival (Relatedness: z = −2.57, df = 1, p = 0.01; Familiarity: z 0 0.34, df = 1, p = 0.73; Relatedness × Familiarity: z = 0.06, df = 1, p = 0.96; Figure 3).

**Figure 3.**
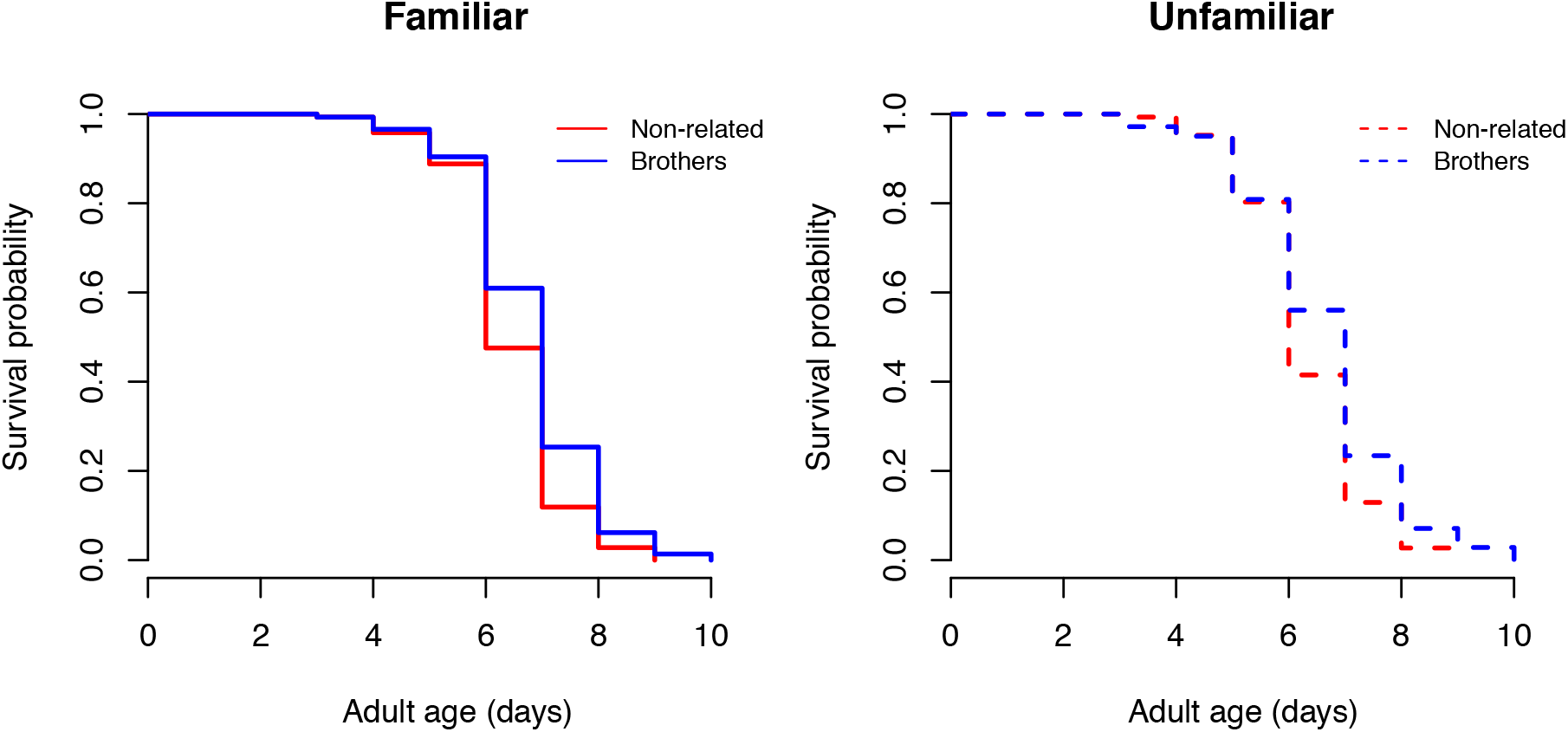
Survival probability for females mated to (A) familiar and (B) unfamiliar males. Blue represents brothers and red represents non-related males.

## 4. DISCUSSION

Adaptive reduction in mate harm can only evolve if 1) there are reliable mechanisms for recognizing kin, and 2) populations are sufficiently viscous (i.e. genetically structured) for relatives to have a reasonable chance of encountering each other while they are reproductively active. The population genetic structure is one challenge facing the hypothesis that kin selection can mitigate the evolution of male harm via interlocus sexual conflict. For example, while kin recognition mechanisms may exist in *Drosophila* (e.g. through cuticular hydrocarbons [45, 46]), Chippindale *et al.* [29] point out that both natural and laboratory populations of *Drosophila* are unlikely to be sufficiently structured to promote kin-selected reduction in male-female conflict. Simply put, adults emerge and fly off and are unlikely to remain in or disperse into genetically structured populations. Le Page *et al.* [32] countered this point by suggesting that genetic structure may occur in fruit flies during colonization of new patches by a small group of females, such that male relatedness-driven reduction of female harm in the established populations are a relic of “the foundation past”. Future work will test whether selection during the foundation of a new population in the natural environment is sufficiently strong to generate long-lasting effects on male reproductive behaviour.

In contrast, *C. maculatus* beetles may, in theory, meet the necessary pre-conditions without difficulty. Female seed beetles lay eggs in clusters, and will deposit all of their eggs in close proximity to each other provided there is sufficient supply of unoccupied beans. Upon emergence from the bean, male seed beetles aggressively court females and begin mating immediately, which increases the probability of encountering relatives. *C. maculatus* is a pest species that infests supplies of stored legumes – under those conditions, it is likely that many beetles will emerge and mate simultaneously providing sufficient variation in the relatedness of competitors. At the same time, we note that *C. maculatus* males are relatively indiscriminate in their mating behaviour, likely because of the high “missing opportunity” cost that is associated with living in high density populations, and commonly mount other males because of perception errors [47, 48], which would complicate selection for fine-tuned kin recognition mechanism that could lead, in theory, to reduced female harm.

In this study, similar to Chippindale *et al.’s* [29] *Drosophila* study, we found that neither male relatedness nor familiarity influenced female lifetime reproductive success. However, in contrast to Chippindale *et al.* [29], male relatedness, but not familiarity, improved female survival. This is also in contrast to Lymbery *et al.* [37], who found that both familiarity and relatedness increase reproductive success, but not survival. In our study, since the effect on survival occurred only in late life, around day six of age when females already stopped producing eggs, it failed to increase female lifetime reproductive success. Therefore, our results do not provide support for the role of kin selection in mitigating the effects of male harm.

Seed beetles are facultatively aphagous – that is, eclosed adults do not require food or water to breed and survive [49]. While in many bruchid beetles adults commonly consume pollen, nectar or fungi [50], *Callosobruchus* beetles do not usually feed as adults. In the current study, we opted to keep the beetles under the aphagous conditions in which they have evolved for over 500 generations since they were first brought into the laboratory in 1979. Thus, our schedule reflects not only the original conditions of the human grain storage, but also the recent evolutionary history of this large outbred population.

Nevertheless, it is interesting to consider how male relatedness could affect the fitness of female beetles in different environments. One may hypothesize that increased survival could result in increased fecundity when beetles have access to additional resources to continue reproduction in late life. Yet, in a recent study by Lymbery *et al.* [37], beetles were raised with the access to yeast but neither lived longer, nor produced more offspring compared to normal non-feeding conditions. This finding seems concordant with earlier reports that did not find positive effects of yeast consumption on fitness in *C. maculatus* [38]. However, despite the lack of positive effects of access to yeast on life-history traits, females kept with related familiar males produced more offspring in Lymbery *et al.* [37] study. This suggests that the presence of yeast interacts with male behaviour in an unknown way to affect female reproductive performance. In our study, we found a three-way interaction between relatedness, familiarity and the shape of age-specific reproduction curve, stemming from the increased early-life reproduction and steeper age-specific decline of females mated to unrelated males then females mated to groups of brothers when both males were raised in a way that precluded familiarity. Increased reproductive performance in early-life could translate into increased individual fitness, but this was not the case. On the other hand, when we included females that failed to produce viable offspring in the analysis, we found that females kept with groups of unrelated males had higher reproductive success, because most of failed reproductive attempts were among females kept with groups of brothers. This finding is in line with the idea that multiple mating increases female fitness when it increases the genetic diversity of partners [51–54].

To summarize, there is little conclusive evidence to date for the role of kin selection in mediating sexual conflict, and, specifically, in reducing male-induced harm to females. More importantly, it is not always easy to see how the selection for such an effect can operate in the natural environment because it requires many opportunities for sib-sib interaction that may not be very common in wild populations of invertebrates [29]. Some organisms, such as bulb mites *(Rhizoglyphus robini),* maybe more prone to the evolution of kin-selected benefits of reduced male harm because of the metapopulation structure characterized by rapid population growth and colonization of the new patches [55]. Indeed, recent work on bulb mites suggest that there is standing genetic variation for male harm that evolves rapidly under kin selection [55], in line with theoretical models [24, 27, 56]. Le Page *et al.* [32] discussed several possible explanations for the fact that reduction in male-induced harm to females is observed in some populations. One likely non-adaptive explanation is a perception error. Indeed, as suggested by Le Page *et al.* [32], males can use cuticular hydrocarbon profiles, or gut microbiota, as a measure of male-male competition, and they may underestimate the level of competition when surrounded exclusively by related males with similar odours, thereby investing less in sperm competition. Such a non-adaptive hypothesis fits squarely with the results of our study, because females housed with groups of related males did enjoy improved survival, suggesting reduced male harm, but this effect was entirely limited to post-reproductive part of their life cycle and had no effect on their individual fitness. We suggest that more work is needed to evaluate the importance of kin selection in the evolution of mating systems.

## Acknowledgements

ERC Starting Grant 2010 AGINGSEXDIFF and ERC Consolidator Grant 2017 GERMLINEAGEINGSOMA to AAM, Swedish Research Council VR grant 2016-05195 to MIL and funding from the American University of Paris to ECB supported this study.

