## Supplementary Figure 1 for "Kin selection and sexual conflict: male relatedness and familiarity do not affect female fitness in seed beetles"

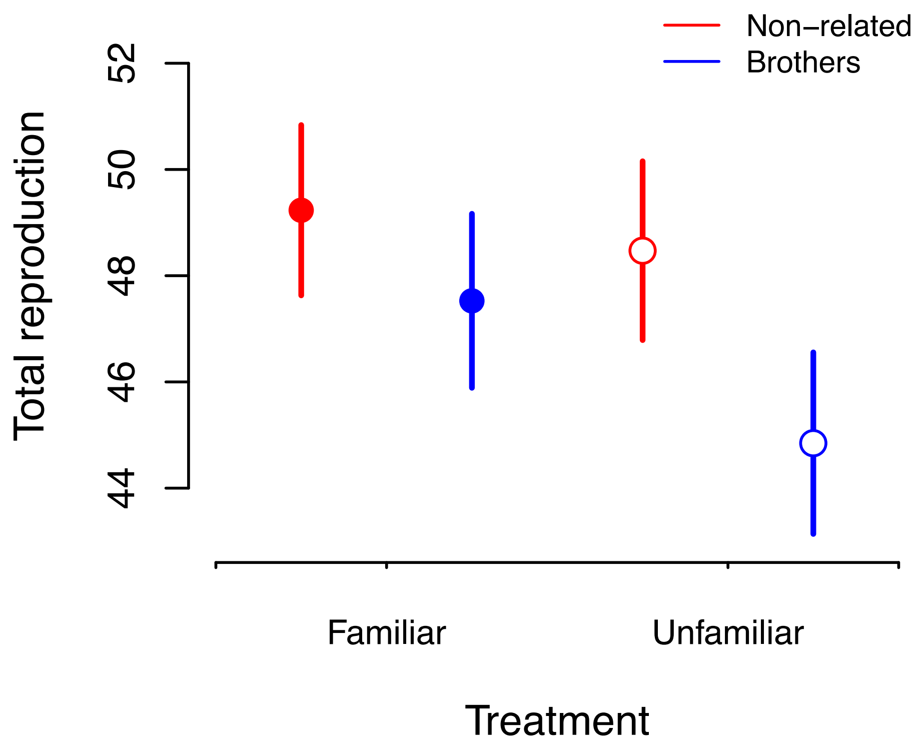


**Supplementary figure 1.** Lifetime reproductive success LRS (number of adult offspring) by treatment group, when including females that did not reproduce: brothers (blue), non-related males (red), familiar individuals (solid symbols) and unfamiliar individuals (open symbols). Symbols represent mean ± SE.
